# Modular capsid decoration boosts adenovirus vaccine-induced humoral and cellular immunity against SARS-CoV-2

**DOI:** 10.1101/2022.02.20.480711

**Authors:** Matthew D. J. Dicks, Louisa M. Rose, Lesley A. H. Bowman, Carl Graham, Katie J. Doores, Michael H. Malim, Simon J. Draper, Mark Howarth, Sumi Biswas

## Abstract

Adenovirus vector vaccines have been widely and successfully deployed in response to COVID-19. However, despite inducing potent T cell immunity, improvement of vaccine-specific antibody responses upon homologous boosting is modest compared to other technologies. Here, we describe a system to enable modular decoration of adenovirus capsid surfaces with protein antigens and demonstrate induction of potent humoral immunity against these displayed antigens. Ligand attachment via a covalent isopeptide bond was achieved in a rapid and spontaneous reaction, requiring simple co-incubation of ligand and vector components. We used a recently described protein superglue, DogTag/DogCatcher, which is similar to the widely used SpyTag/SpyCatcher ligation system but performs better in loop structures. DogTag was inserted into surface-exposed loops in the adenovirus hexon protein to allow attachment of DogCatcher-fused ligands on virus particles. Efficient coverage of the capsid surface was achieved using a variety of ligands and vector infectivity was retained in each case. Capsid decoration shielded particles from anti-adenovirus neutralizing antibodies. In prime-boost regimens, proof-of-concept COVID-19 adenovirus vaccines decorated with the receptor-binding domain (RBD) of SARS-CoV-2 spike induced >10-fold higher SARS-CoV-2 neutralization titers compared to an undecorated adenovirus vector encoding spike. Importantly, decorated vectors retained robust T cell immunogenicity to encoded antigens, a key hallmark of adenovirus vector vaccines. We propose capsid decoration via protein superglue-mediated covalent ligation as a novel strategy to improve the efficacy and boostability of adenovirus-based vaccines and therapeutics.

**One Sentence Summary:** Decorating the capsid surface of adenovirus vaccine vectors using a spontaneous protein superglue induces potent pathogen-specific immunity

## INTRODUCTION

The COVID-19 pandemic has demonstrated the remarkable utility of replication-defective recombinant adenoviruses as vaccine vectors (*1*). The rapid construction, scalability and cost-effectiveness of this platform, without a requirement for adjuvants, combined with long-term stability at standard refrigerator temperatures, has positioned the platform among leading technologies in the pandemic response (*1*). There are currently four adenovirus-based COVID-19 vaccines licensed and in use across various parts of the world, using a range of adenovirus serotypes (*2-5*).

However, despite decades of research and significant progress in the development of this vaccine platform, limitations remain. While adenovirus vectors are among the most potent inducers of cellular immunity (particularly CD8^+^ T cell responses (*6*)) in humans, antibody responses to target antigens are modest in comparison to the most potent inducers of humoral immunity, which include nanoparticle and virus-like-particle (VLP) based recombinant protein technologies (*7, 8*). Anti-vector immunity (particularly via anti-capsid neutralizing antibodies) has also been shown to limit vaccine immunogenicity and efficacy, since transduction of host cells by the viral vector prior to immune clearance is essential to induce immunity against encoded target antigens (*5, 9, 10*). Human populations harbor significant pre-existing neutralizing antibody titers against common human adenovirus serotypes, which has led to a search for adenovirus serotypes with lower human seroprevalence (including Ad26 (*11*) and Y25/ChAdOx1 (*12*)). However, with the most clinically advanced serotypes already used to vaccinate more than a billion people during the current pandemic, it may be challenging to re-use these platforms for boosting and subsequent disease indications. Previous studies have shown that the efficacy of prime-boost regimens using the same vector for both immunizations is limited by the anti-vector neutralizing responses raised after the first immunization (*9*).

To address these limitations, here we have designed a platform to enable modular covalent decoration of adenovirus particles with antigenic ligands. High-density repetitive display of antigen on virus-like-particles (VLPs) is a highly effective strategy to generate potent and boostable humoral immunity (*13*). The SpyTag/SpyCatcher protein superglue system has previously been utilized to achieve rapid and efficient covalent attachment of vaccine antigens to a variety of VLP platforms (*14-17*). SpyTag, a short peptide tag, reacts rapidly and spontaneously with SpyCatcher, a small protein domain, forming an irreversible isopeptide bond (*18*). A hepatitis-B surface antigen (HBsAg) VLP-based COVID-19 vaccine candidate displaying the receptor-binding domain of SARS-CoV-2 spike (RBD) using SpyTag/SpyCatcher technology has recently entered Phase I/II clinical trials (*19*). Protein superglue technologies can overcome challenges associated with other methods of antigen attachment to VLPs, such as irregular distribution and orientation of ligands coupled using chemical modification, or protein/particle instability following genetic fusion/insertion of ligands to particle scaffolds (*20*). Recently, a new protein superglue pair based on domain 4 of the RrgA adhesin from *Streptococcus pneumoniae*, DogTag/DogCatcher, was shown to perform particularly efficiently in loop structures (Fig. S1)(*21*).

Here, we used the DogTag/DogCatcher system to enable modular capsid display of protein ligands including RBD on the surface of recombinant adenovirus vectors. DogTag was engineered into surface loops on the hexon capsid protein and enabled attachment of DogCatcher-fused antigens. Despite extensive coverage of the capsid surface, decorated virions retained infectivity and induced potent neutralizing antibody and T cell responses against SARS-CoV-2 in a proof-of-concept study. Capsid display markedly improved boostability of responses compared to homologous prime-boost regimens using conventional undecorated adenovirus vaccines. To the best of our knowledge, this is the first example of covalent decoration of adenovirus vector capsids using spontaneous isopeptide linkage and represents a new tool for the development of adenovirus-based vaccines and therapeutics.

## RESULTS

### Adenovirus capsid decoration through spontaneous isopeptide bond linkage

The hexon protein is the major component of the adenovirus capsid (each virion comprising 720 copies assembled into 240 trimers (*22*)) and is therefore an ideal target for modular capsid display of antigens (Fig. 1A). We utilized the DogTag/DogCatcher reactive pair to achieve spontaneous covalent coupling of DogCatcher-fused ligands onto the surface of recombinant adenovirus vectors. To enable covalent capsid decoration, DogTag (23 amino acids) was genetically inserted into each of three hexon hypervariable surface loops (HVR1, HVR2 or HVR5) flanked by flexible glycine-serine linker sequences (Fig. 1B). Replication-defective (E1-deleted) recombinant adenovirus vectors with DogTag insertions (Ad-DogTag) were readily produced in E1 complementing 293A cells, with no reduction in yield compared to similar vectors with an unmodified hexon (Fig. 2A and Fig. S2). Vectors with SpyTag inserted at each of the hexon HVRs were also produced at equivalent titers (Fig. S2). Particle to infectious unit ratios (P:I ratios) of vector preparations were comparable between vectors with and without hexon insertions (Fig. 2A and Fig. S2). Co-incubation of Ad-DogTag virions with recombinant DogCatcher protein achieved >90% coverage of the Ad capsid (>90% hexon protein covalently coupled to DogCatcher as measured by SDS-PAGE gel shift, Fig. 2B). Similar levels of coverage were observed with DogTag inserted at each of the three hexon HVRs, indicating a high degree of flexibility in this approach. Strikingly, despite extensive coverage of the capsid with DogCatcher, decorated virions with DogTag at HVR2 or HVR5 exhibited no reduction in infectivity (<1.5-fold) *in vitro* compared to undecorated virions (Fig. 2C). Decorated virions with DogTag at HVR1 exhibited a modest 1.7-fold reduction (Fig. 2C). In contrast, SpyTag was poorly reactive with SpyCatcher when inserted into Ad hexon HVRs, with markedly lower capsid coverage compared to the DogTag/DogCatcher system, despite co-incubation with a 7-fold higher concentration of recombinant SpyCatcher protein (Fig. S3A). Furthermore, once significant capsid coverage with SpyCatcher was achieved (i.e. 57% hexon protein coupled), virion infectivity *in vitro* was reduced ∼100-fold (Fig. S3B).

**Fig. 1.**
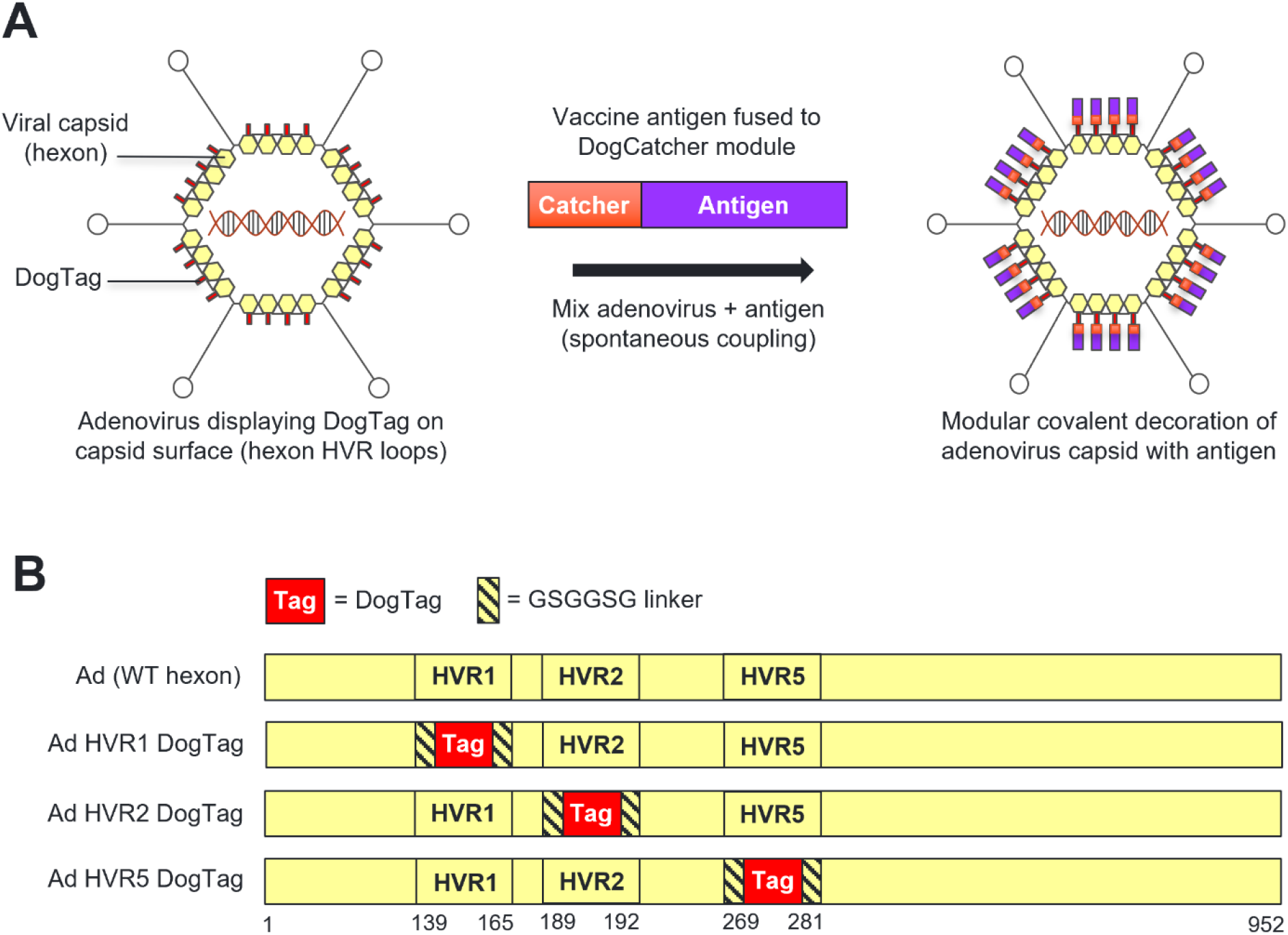
Modular covalent decoration of the adenovirus capsid via insertion of DogTag into hexon HVR loops. (**A**) Modular display of DogCatcher-fused antigenic ligands on the surface of the adenovirus capsid via covalent coupling with DogTag inserted into hexon HVR surface loops. Attachment of antigens to the capsid achieved by simple co-incubation of adenovirus and antigen components in a rapid and spontaneous reaction. (**B**) Design of modified adenovirus hexon sequences with DogTag inserted into either HVR1, HVR2 or HVR5 flanked by flexible linkers. Amino acid residue numbers corresponding to deletion / insertion sites at HVR1, HVR2 or HVR5 in Ad5 hexon are indicated.

**Fig. 2.**
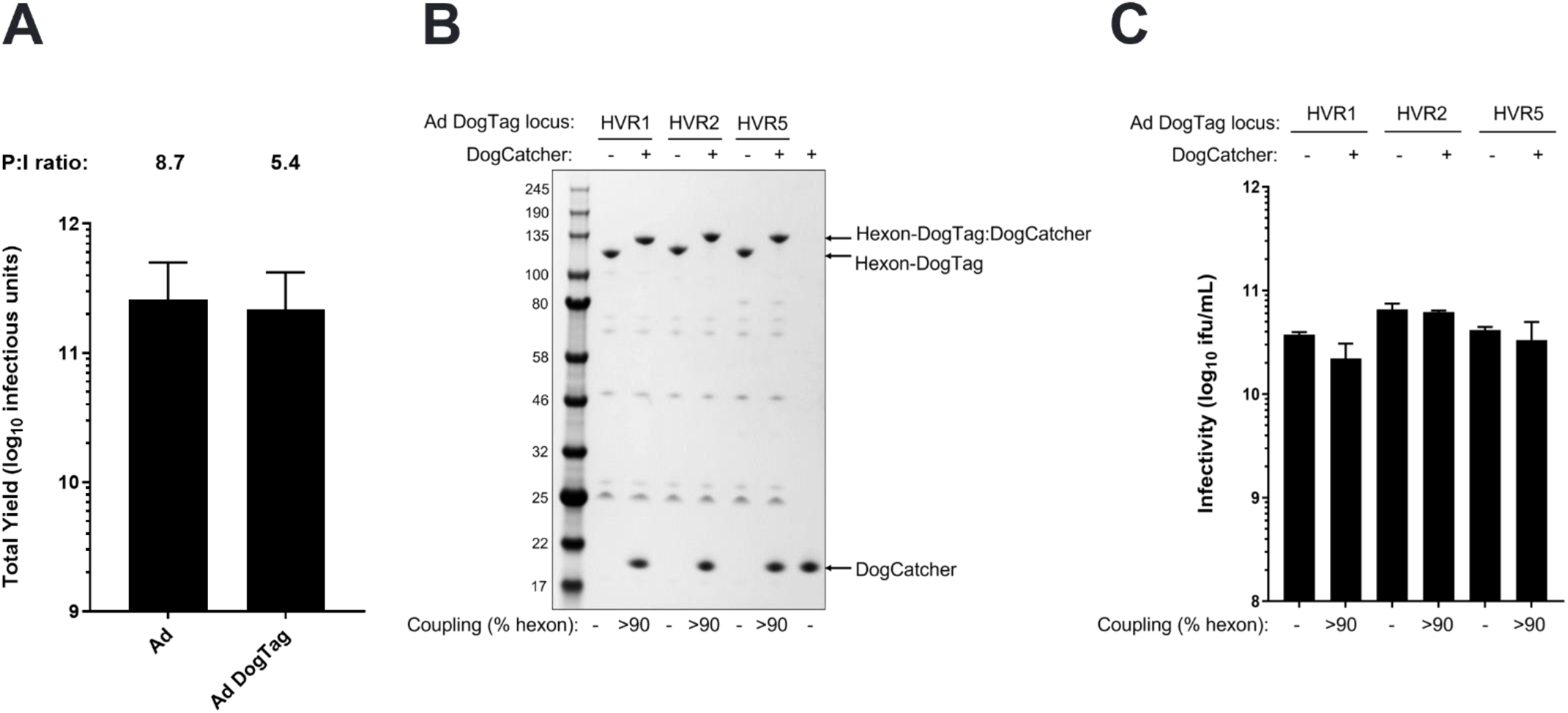
DogTag is highly reactive with DogCatcher following insertion into adenovirus hexon HVR loops, with decorated virions retaining infectivity. (**A**) Yield comparison of GFP-expressing Ad vector preparations displaying DogTag on hexon HVR5 (Ad DogTag) versus Ad vectors with an unmodified hexon (Ad). Data show mean + SD, n=3, infectious yield from 1500 cm^2^ adherent 293A cells. Mean P:I ratios (ratio of total viral particles calculated by UV spectrophotometry to infectious units calculated by GFP focus assay) for vector batches are indicated above each bar. (**B**) SDS-PAGE and Coomassie staining analysis of Ad virions displaying DogTag at HVR1, HVR2 or HVR5 (1E+10 viral particles) incubated with DogCatcher (5 μM) at 4°C for 16 h. Gel shift observed upon covalent coupling of DogCatcher to virion associated hexon-DogTag. (**C**) Vector infectivity (GFP focus assay) performed in 293A cells on the samples from B. Data show mean + SD of triplicate wells.

### Decoration of virions with a model antigen elicits potent humoral immunity

The circumsporozoite protein (CSP) of *Plasmodium falciparum* (Pf) has been extensively studied as a malaria vaccine candidate antigen (*23*). The protein contains a highly immunogenic repetitive region, primarily consisting of repeats of the amino acid sequence NANP; vaccine-induced antibodies against this NANP repeat region protect against malaria infection (the licensed malaria vaccine RTS,S/Mosquirix™ includes 18 copies of NANP (*23*)). A model system to test antibody induction using capsid display technology was developed by fusing multiple copies of NANP (either 9 or 18) to DogCatcher. These DogCatcher-NANP_n_ fusion proteins were used to decorate Ad-DogTag virions. Extensive coverage of the Ad capsid (>90% hexon coupled) was achieved using both DogCatcher-NANP_n_ proteins (Fig. 3A). Neither ligand had any effect on vector infectivity upon particle decoration (Fig. 3B). Since coverage of Ad virions was extensive with both ligands, we sought to determine whether particle decoration could act as a protective shield to inhibit Ad neutralization by capsid-binding antibodies. Decoration with both ligands partially shielded Ad particles from neutralization *in vitro* by an anti-hexon monoclonal antibody (mAb 9C12 (*24*)), increasing the IC_50_ by at least 10-fold compared to Ad-DogTag alone (Fig. 3C). Particle decoration also inhibited neutralization of Ad particles by polyclonal anti-Ad5 serum (Fig. 3D), even though antibodies in this serum were raised against multiple capsid components. Co-incubation of Ad-DogTag virions with SpyCatcher (SpyCatcher does not react with DogTag, hence no capsid decoration (*21*)) did not inhibit neutralization with anti-Ad5 sera (Fig. 3D).

**Fig. 3.**
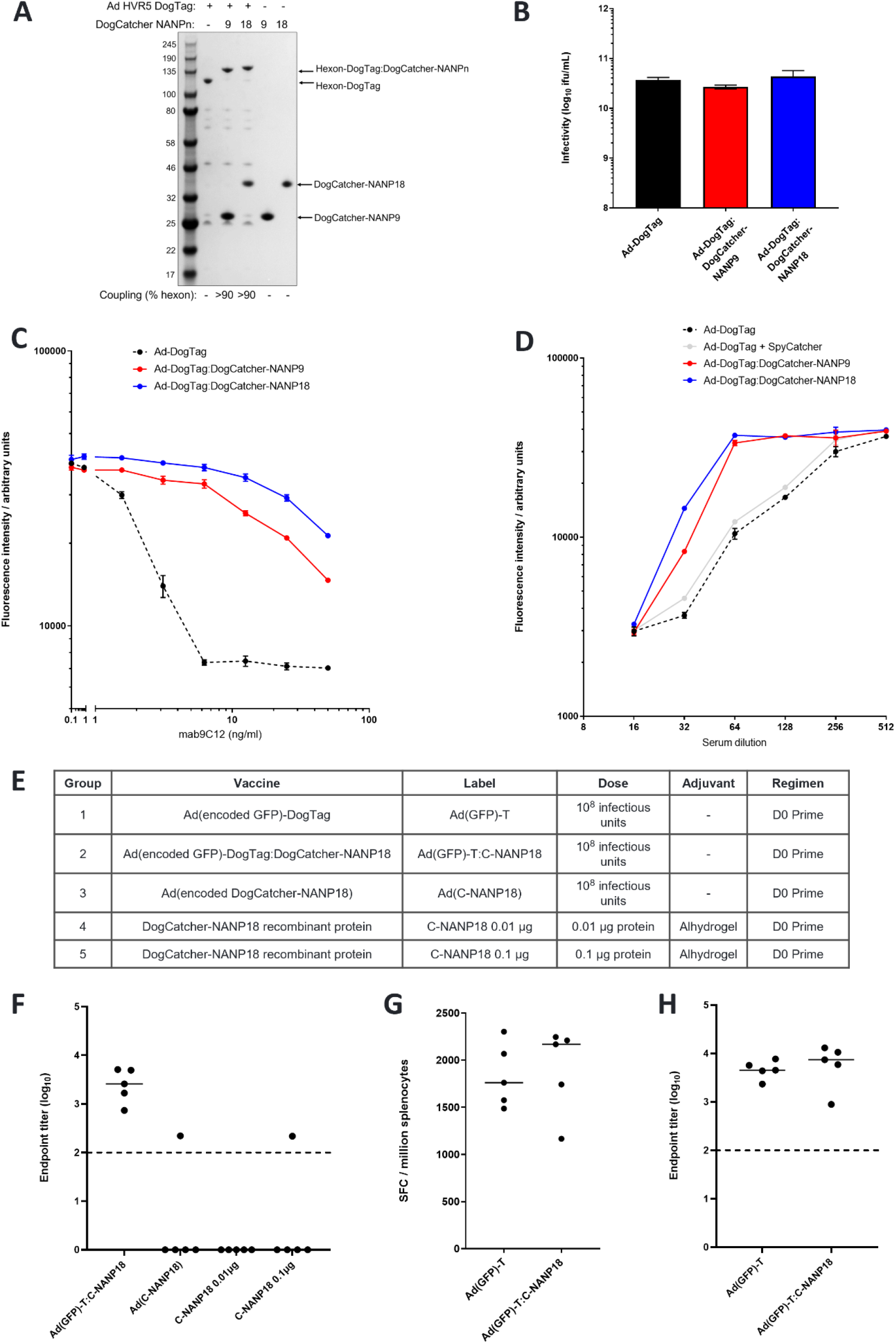
Capsid display of DogCatcher-NANP_18_ ligand shields the particles from anti-vector neutralizing antibodies and elicits potent ligand-specific humoral immunity. (**A**) SDS-PAGE and Coomassie staining analysis of Ad virions displaying DogTag at HVR5, (1E+10 viral particles) incubated with DogCatcher-NANP_9_ (5 μM) or DogCatcher-NANP_18_ (5 μM) at 4°C for 16 h. (**B**) Vector infectivity (GFP focus assay) performed in 293A cells on the samples from A. Data show mean + SD of triplicate wells. (**C**) Vector neutralization assay using anti-hexon mAb 9C12. Ad particles encoding a GFP transgene with or without a capsid ligand were added to 293A cells in the presence of a varying concentration of neutralizing mAb. Vector transduction was measured via fluorescence of encoded GFP expressed in the cells. (**D**) Vector neutralization assay using anti-Ad5 serum. Ad particles encoding a GFP transgene with or without a capsid ligand were added to 293A cells in the presence of a varying concentration of serum from mice immunized with Ad5. C-D show mean + range of duplicate wells. (**E**) BALB/c mice (5 per group) were immunized intramuscularly as described (vector encoded antigens in brackets). The DogCatcher-NANP18 protein dose in Group 2 was calculated to be <0.05 μg per mouse. (**F**) Serum IgG antibody responses to DogCatcher-NANP18 in Groups 2-5 measured by endpoint ELISA 14 days post-immunization. (**G**) CD8^+^ T cell responses in the spleen to encoded epitope EGFP_200-208_ were measured in Groups 1 and 2 by overnight *ex vivo* IFNγ-ELISpot 14 days post-immunization. SFC = spot forming cells. (**H**) Serum IgG antibody responses to encoded GFP in Groups 1 and 2 were measured by endpoint ELISA 14 days post-immunization. In F-H, bars show median responses. Dashed line represents limit of detection.

An immunogenicity study in mice was performed to assess antibody titers against DogCatcher-NANP_18_ displayed on Ad-DogTag virions, as compared to a conventional Ad vector encoding the same model antigen construct or vaccination with protein in adjuvant formulations (Fig. 3E, Groups 2-5). Two weeks after a single vaccine dose, robust IgG antibody responses against DogCatcher-NANP_18_ were generated using capsid display technology (Ad(GFP)-DogTag:DogCatcher-NANP18), but were not achieved using an encoded antigen platform (Ad(DogCatcher-NANP18) or protein in adjuvant formulations (DogCatcher-NANP18) (Fig. 3F). In the same experiment, we also assessed the impact of capsid decoration of an Ad particle on immune responses generated against a transgene-encoded antigen in the same vector (Fig. 3E, Groups 1 and 2). No differences in magnitudes of either CD8^+^ T cell responses (Fig. 3G) or antibody responses (Fig. 3H) against encoded GFP were observed between Ad-DogTag vectors with capsid decoration (Ad(GFP)-DogTag:DogCatcher-NANP18) or without capsid decoration (Ad(GFP)-DogTag). These data demonstrate that, despite >90% hexon coverage on Ad virions, display of DogCatcher-NANP_18_ ligand on the capsid surface did not impair immune responses against an encoded transgene antigen.

### Decoration with SARS-CoV-2 spike receptor-binding domain (RBD) shields adenovirus particles from capsid interactors

The receptor-binding domain (RBD) of SARS-CoV-2 spike protein interacts with the angiotensin-converting enzyme 2 (ACE2) on susceptible host cells and is required for viral infection. A recent study has estimated that ∼90% of SARS-CoV-2 neutralizing antibodies raised against the spike protein target the RBD region (*25*). As further proof-of-concept for Ad capsid display technology, we fused RBD (∼26kDa) to DogCatcher (Fig. 4A), and decorated Ad-DogTag virions with DogCatcher-RBD fusion protein. Despite the larger size of DogCatcher-RBD compared to ligands tested previously, substantial coverage of the Ad capsid was achieved (∼57% hexon monomers coupled) (Fig. 4B), and importantly no impairment of vector infectivity was observed upon particle decoration (Fig. 4C). Strikingly, capsid display of DogCatcher-RBD completely abrogated Ad vector neutralization *in vitro* by potent hexon-binding neutralizing monoclonal antibody mAb 9C12 (Fig. 4D). Vector neutralization by polyclonal anti-Ad5 serum was also inhibited by capsid shielding with DogCatcher-RBD (Fig. 4E). Given the observed capacity for DogCatcher-RBD to shield the Ad capsid from antibody-mediated neutralization, we sought to determine whether decoration with this ligand could impair other Ad capsid interactions. Human coagulation factor X (hFX) has previously been shown to bind directly to the Ad5 hexon protein and mediates hepatocyte transduction via heparan sulfate proteoglycans (HSPGs) by Ad5 vectors *in vivo*, particularly after intravascular delivery (*26, 27*). Using a cell-line model of this interaction, capsid decoration with DogCatcher-RBD completely abrogated hFX-mediated Ad-DogTag transduction of SKOV-3 cells. Ad-DogTag without a capsid shield exhibited a 6.6-fold increase in SKOV-3 infectivity upon co-incubation with hFX (Fig. 4F). Decorated Ad virions exhibited a modest reduction in infectivity (1.8-fold) in the absence of hFX compared to undecorated virions (Fig. 4F).

**Fig. 4.**
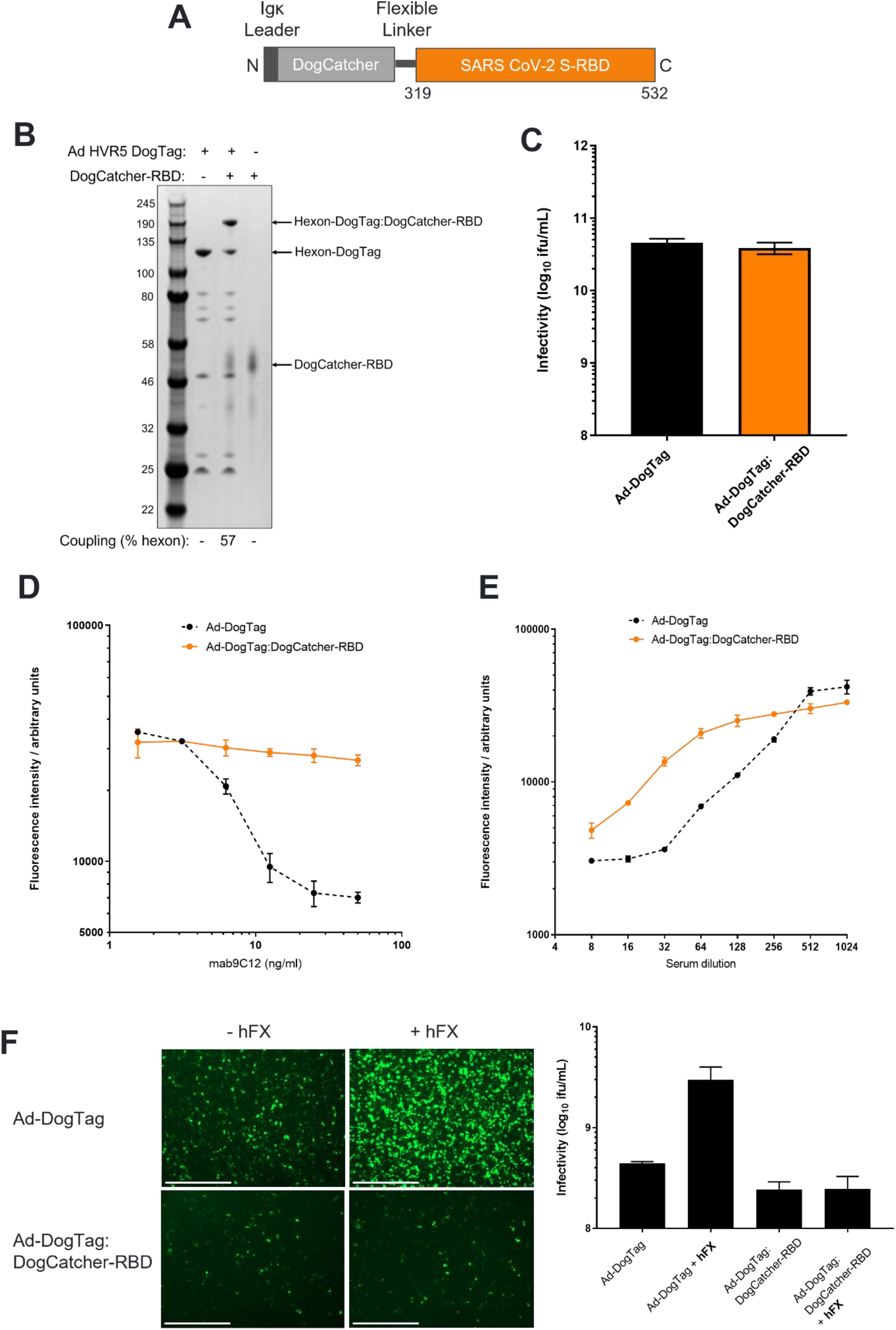
RBD can be fused to DogCatcher, displayed on adenovirus particles, and is an effective capsid shield. (**A**) Design of DogCatcher-RBD protein (**B**) SDS-PAGE and Coomassie staining analysis of Ad virions displaying DogTag at HVR5 (Ad-T) (1E+10 viral particles) incubated with DogCatcher-RBD (3.5 μM) at 4°C for 16 h. (**C**) Vector infectivity (GFP focus assay) performed on the samples from B. Data show mean + SD of triplicate wells. (**D**) Vector neutralization assay using anti-hexon mAb 9C12, performed as described in Fig. 3. (**E**) Vector neutralization assay using anti-Ad5 sera, performed as described in Fig. 3. C-D show mean + range of duplicate wells. (**F**) Capsid decoration with DogCatcher-RBD impairs human Factor X (hFX)-mediated Ad transduction of SKOV3 cells. Fluorescent microscopy images taken 48 h post-infection show Ad-DogTag or Ad-DogTag:DogCatcher-RBD vectors expressing GFP incubated with or without hFX (2500 viral particles / cell). Scale bar = 1000 µm. Infectivity data (GFP focus assay) show mean + SD of triplicate wells.

### CryoEM analysis of adenovirus particles displaying RBD

To further investigate the nature of capsid decoration of Ad-DogTag particles with DogCatcher-RBD, cryoEM analysis of decorated (Ad-DogTag:DogCatcher-RBD) and undecorated (Ad-DogTag) particles was performed (Fig. 5). A 3D density map of decorated particles clearly indicated additional density extending outward from hexon trimers (red-orange regions), as compared to the undecorated particles (Fig. 5A). Additional density protruding from hexon trimers could also be observed from 2D class averages of Ad-DogTag:DogCatcher-RBD particles compared to Ad-DogTag particles, extending the particle diameter from 93 nm to 98 nm (Fig. 5B). Surface protrusions (representing hexon trimers) on Ad-DogTag were of uniform density, while Ad-DogTag:DogCatcher-RBD showed two distinct regions. The outermost region represented coupled RBD being less dense than hexon trimers, likely due to flexibility of the attached ligand. Analysis of 3D density maps revealed attachment of either one (type I, Fig. 5C) or two (type II, Fig. 5D) RBD ligands per hexon trimer. Type II decoration was particularly abundant on hexon trimers located adjacent to penton subunits at capsid vertices. These trimers are elevated with respect to the rest of the virion surface (see Fig. 5A, red-orange regions of Ad-DogTag map; note that the fiber protein is not shown) and therefore steric hindrance may be lower, enabling multiple ligand occupancy. Additional density representing attached RBD is clear for both type I and type II arrangement, as compared to undecorated Ad-DogTag; Fig. S4 shows fitting of an RBD structure into these maps (with the RBD N-terminus proximal to hexon HVR5, DogCatcher is not shown).

**Fig. 5.**
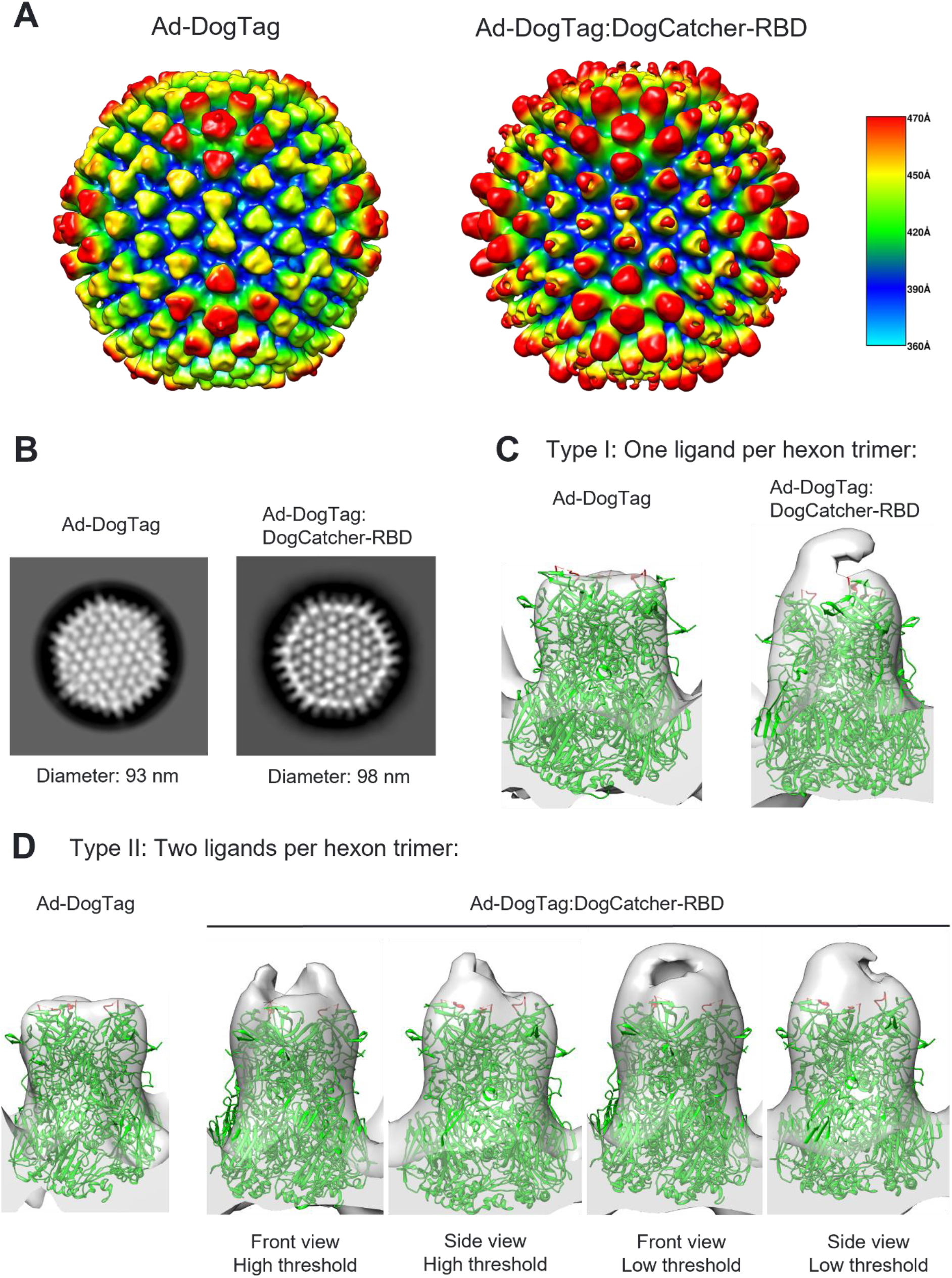
CryoEM analysis of adenovirus particles displaying RBD. (**A**) 3D density maps (at 10.5 Å) for Ad-DogTag (control sample, undecorated) and Ad-DogTag:DogCatcher-RBD particles. Radial coloring scheme is indicated; regions furthest from the center of the particle are shown in red, regions closest to the center are shown in blue. Distance from center of particle (in angstroms) is indicated. (**B**) Exemplary 2D class averages. Indicated diameters calculated from vertex to vertex. (**C**) Type I ligand coupling; 3D structure of representative hexon trimer without ligand (Ad-DogTag) or with one ligand coupled per trimer (Ad-DogTag:DogCatcher-RBD) shown at the same contour level. (**D**) Type II ligand coupling; 3D structure representative of hexon trimer adjacent to penton base without ligand (Ad-DogTag) or with two ligands coupled per trimer (Ad-DogTag:DogCatcher-RBD) shown at the same contour level). Maps for Ad-DogTag:DogCatcher-RBD shown at both front and side angles, and high and low threshold to indicate location and extent of additional electron density. In both C and D, hexon trimer structure (PDB ID 6B1T) was fitted (green), with location of HVR5 loop (residues 270-280, site of DogTag insertion) shown in red.

### Capsid decoration with RBD elicits potent neutralizing antibody responses against SARS-CoV-2

An *in vivo* study was conducted to assess humoral and cellular immunity against SARS-CoV-2 using Ad-DogTag vectors decorated with DogCatcher-RBD, compared to a conventional adenovirus vector encoding spike in the viral genome.

Mice were immunized in homologous prime-boost regimens with either Ad-DogTag encoding spike without a capsid ligand (Ad(Spike)-DogTag, Group 1), the same vector as in Group 1 but with DogCatcher-RBD coupled to the viral capsid (Ad(Spike)-DogTag:DogCatcher-RBD, Group 2), a vector displaying the same DogCatcher-RBD ligand as in Group 2 but encoding a GFP transgene rather than spike (Ad(GFP)-DogTag:DogCatcher-RBD, Group 3), or recombinant DogCatcher-RBD protein in Alhydrogel adjuvant (Group 4) (Fig. 6A).

**Fig. 6.**
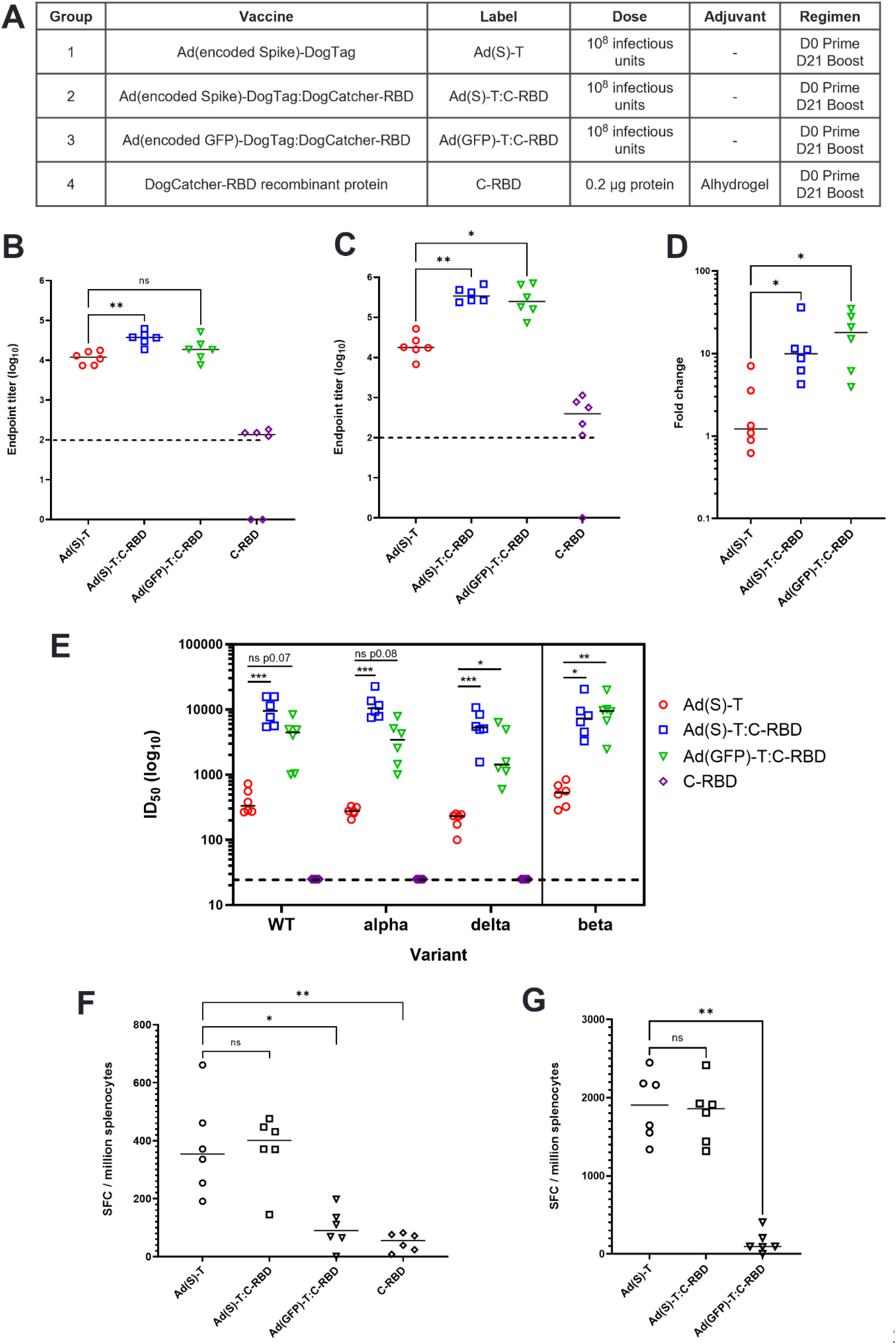
High-titer SARS-CoV-2 neutralizing antibody responses generated by capsid display of RBD. (**A**) BALB/c mice (6 per group) were immunized intramuscularly in homologous prime-boost regimens as described. The DogCatcher-RBD protein dose in Groups 2 and 3 was calculated to be <0.2 μg per mouse. (**B**) Serum IgG antibody responses to RBD at D20 measured by endpoint ELISA. (**C**) Serum IgG antibody responses to RBD at D35 measured by endpoint ELISA. (**D**) Fold increase in anti-RBD IgG titer post-boost (D35 titer in C divided by D20 titer in B) (**E**) SARS-CoV-2 neutralization titers (pVNT assay) in D35 serum against Wuhan strain (WT) and variants of concern alpha (B.1.1.7), delta (B.1.617.2) and beta (B.1.351). Data on beta variant collected separately. (**F**) IFNγ-ELISpot response in spleen at D35 against peptide pool spanning SARS-CoV-2 S-RBD. (**G**) IFNγ-ELISpot response in spleen at D35 against peptide pool spanning full length SARS-CoV-2 S protein. Dashed lines represent limit of detection. Median responses shown by a horizontal line.

Serum IgG antibody titers against RBD were comparable between Ad vector groups after a single immunization (Fig. 6B), but median post-boost titers were >10-fold higher using both RBD capsid display vectors compared to the conventional Ad(Spike)-DogTag vector, regardless of the encoded antigen (Fig. 6C). Indeed, anti-RBD IgG titers after immunization (both pre-and post-boost) with Ad(GFP)-DogTag:DogCatcher-RBD were comparable to Ad(Spike)-DogTag:DogCatcher-RBD, despite the former vector lacking an encoded spike antigen. A comparison of median anti-RBD IgG responses pre-and post-boost indicates that capsid display of RBD increased boostability of Ad vector mediated humoral immunity against RBD ∼10-fold (Fig. 6D). IgG responses against the full-length spike ectodomain were also higher after immunization with both RBD capsid display vectors post-boost (Fig. S5). Neutralizing antibody titers in serum post vaccination were measured in an extensively validated *in vitro* neutralization assay using human immunodeficiency virus-1 (HIV-1) based virus particles pseudotyped with SARS-CoV-2 spike (*28*). Median pseudotype neutralization titers (pVNT) after prime-boost immunization with either RBD capsid display vector were >10-fold higher against the Wuhan strain compared to the conventional Ad vector encoding spike (Fig. 6E). Similar relative titers between groups were observed against alpha, beta and delta variants of concern (Fig. 6E).

Importantly and in agreement with previous observations, display of DogCatcher-RBD on the Ad capsid surface did not impair T cell responses against encoded SARS-CoV-2 spike. Spleen interferon-gamma (IFNγ) enzyme-linked immune absorbent spot (ELISpot) responses against an RBD peptide pool (Fig. 6F) and a full-length spike peptide pool (Fig. 6G) showed that CD8^+^ T cell immunogenicity with Ad(Spike)-DogTag and Ad(Spike)-DogTag:DogCatcher-RBD was comparable. CD8^+^ T cell responses against RBD were significantly higher in magnitude after immunization with either Ad vector encoding spike, compared to Ad(GFP)-DogTag:DogCatcher-RBD (Dunn’s multiple comparisons test; Ad(Spike)-DogTag vs Ad(GFP)-DogTag:DogCatcher-RBD p=0.015, Ad(Spike)-DogTag:DogCatcher-RBD vs Ad(GFP)-DogTag:DogCatcher-RBD p=0.009), indicating that encoding an antigen is preferable to capsid display for inducing CD8^+^ T cell responses.

### Applying a capsid shield to booster adenovirus vaccines improves both humoral and cellular immunogenicity

Given the ability of capsid display technology to elicit potent humoral immunity against SARS-CoV-2, we sought to determine whether RBD capsid decoration could efficiently boost immunity in animals that had already received a first dose of a conventional Ad vector encoding spike. A homologous two-dose Ad(Spike)-DogTag regimen was compared to a heterologous regimen in which an RBD capsid shield (Ad(Spike)-DogTag:DogCatcher-RBD) was applied to the second vaccine dose (Fig. 7A). A 5.3-fold increase in median anti-RBD IgG titers was observed between prime and boost immunizations in the homologous regimen, but a 63.1-fold increase in titer was observed after the heterologous regimen (Fig. 7B). In addition, spleen IFNγ ELISpot responses against full-length SARS-CoV-2 spike were also higher in the heterologous regimen compared to the homologous regimen (Fig. 7C). A similar observation was observed in IFNγ ELISpot assays using a peptide pool spanning only residues 633-1273 of spike (hence not including the RBD region), indicating that increased cellular immunity to the encoded spike transgene after heterologous prime-boost occurred independently from the contribution of the RBD capsid ligand (Fig. 7D).

**Fig. 7.**
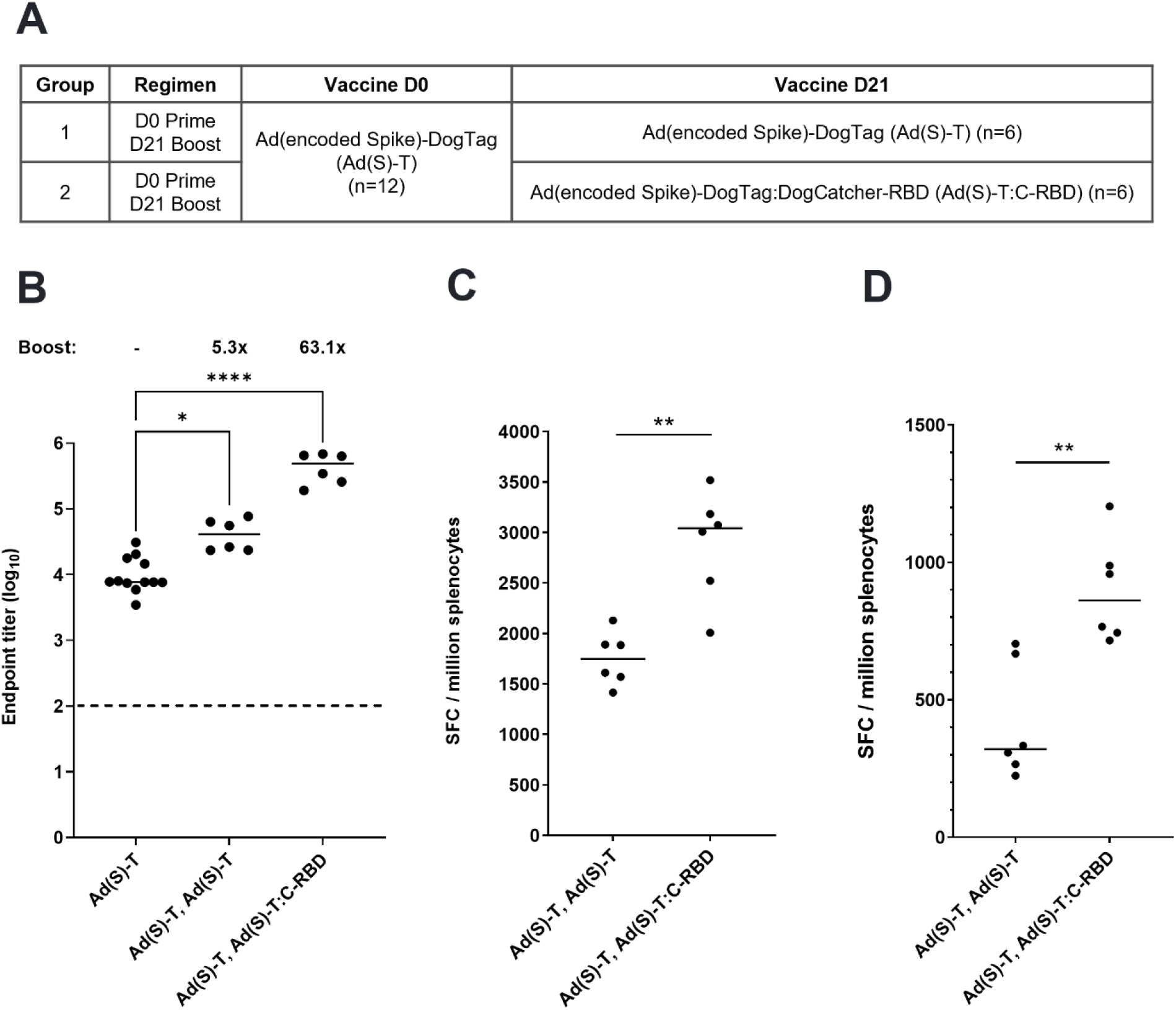
Applying RBD capsid decoration at boost significantly increases SARS-CoV-2 specific antibody and T cell responses in an adenovirus vector prime-boost regimen. (**A**) BALB/c mice (n=12) were immunized intramuscularly with Ad(Spike)-DogTag on D0 and then on D21 given a second intramuscular immunization of either Ad(Spike)-DogTag (Group 1, n=6) or Ad(Spike)-DogTag:DogCatcher-RBD (Group 2, n=6). All vaccines were administered at a dose of 10^8^ infectious units. (**B**) Serum IgG antibody responses to RBD measured by endpoint ELISA. Responses measured post prime on D20 (Ad(Spike)-DogTag) are compared to responses on D35 after homologous (Ad(Spike)-DogTag, Ad(Spike)-DogTag) or heterologous (Ad(Spike)-DogTag, Ad(Spike)-DogTag:DogCatcher-RBD) prime boost. Fold change in median titer post boost displayed. Dashed line represents limit of detection. (**C**) IFNγ-ELISpot response in spleen at D35 against peptide pool spanning full length (1-1273) SARS-CoV-2 S. (**D**) IFNγ-ELISpot response in spleen at D35 against peptide pool spanning C-terminal residues 633-1273 of SARS-CoV-2 S only (i.e. not including RBD domain). In B-D, horizontal bars show median responses. The dashed line represents the limit of detection.

## DISCUSSION

In this study we describe a simple, rapid, and targeted system of spontaneous covalent decoration of recombinant adenovirus particles with a variety of ligands under mild conditions. Using the recently described DogTag/DogCatcher reactive pair (*21*), we targeted the hexon protein for capsid display to maximize ligand density since hexon is the most abundant component of the adenovirus capsid. Several previous studies have achieved display of short immunogenic epitopes by direct genetic insertion of sequences into hexon HVR loops. However, these approaches have typically been limited to <50 residues, since larger insertions disrupt protein stability and prevent successful rescue of recombinant vectors (*29-31*). Other studies have performed chemical modification of the viral capsid to attach larger ligands, including targeted insertion of cysteine residues for thiol-directed coupling, but these approaches can require challenging reaction and storage conditions to achieve conjugation and retain infectivity of modified vectors (*32, 33*). Some studies achieved capsid display of larger ligands through genetic fusion to the C-terminus of pIX, a minor capsid protein with fewer copies per virion than hexon (*30*). However, large pIX fusions do not always generate viable vectors, can reduce capsid stability, and modified pIX may not be efficiently incorporated into the capsid (*34, 35*).

Using our approach, we have covalently coupled ligands as large as the RBD of SARS-CoV-2 spike (∼26 kDa) to hexon through genetic fusion to DogCatcher and co-incubation of the recombinant fusion protein with Ad-DogTag, achieving extensive coverage of the viral capsid in each case (Fig. 2B, 3B, 4B and 5A). Importantly and despite a high proportion of hexon monomers conjugated to ligand, decorated vectors retained infectivity *in vitro* (Fig. 2C, 3C and 4C). Transduction of 293A cells by Ad5 vectors is mediated by interactions between the adenovirus fiber protein and the coxsackie and adenovirus receptor (CAR)(*36*); these interactions are unlikely to be masked by hexon conjugation due to the sizes of the ligands tested, as reflected by our cryoEM data (Fig 5). We hypothesize that there is also sufficient flexibility engineered into hexon loop structures to avoid interference with secondary interactions between the adenovirus penton and cell-surface αV integrins required for clathrin-mediated endocytosis of virions (*37*). Mechanisms of Ad vector transduction upon *in vivo* delivery may be more complex than those described *in vitro* (*38*). However, since transduction of cells is an essential prerequisite for expression of encoded transgenes and neither humoral nor cellular transgene-specific immune responses are impaired by capsid decoration (Fig. 3 and Fig. 6), our data suggest that decorated Ads also efficiently transduce cells *in vivo* (though we have not investigated this directly). DogTag was highly reactive with DogCatcher at each of the three insertion sites tested (Fig. 2B), with decorated vectors retaining infectivity in each case (Fig. 2C). These data indicate that reactivity is not insertion site-specific and imply that our approach is likely to be successful with alternative adenovirus serotypes.

Decoration of the Ad capsid surface with DogCatcher-RBD reduced the potency of anti-Ad neutralizing antibodies *in vitro*. Capsid shielding against mAb 9C12 was more effective with DogCatcher-RBD (Fig. 4D) than DogCatcher-NANP18 (Fig. 3C), despite lower ligand coverage (57% (Fig. 4B) versus >90% (Fig. 3A) hexon coupled), likely due to increased coverage of the capsid surface with the larger RBD ligand. Indeed, cryoEM data indicated that all hexon trimers had at least one copy of RBD attached, with ligand density covering much of the hexon trimer surface (Fig. 5). Shielding from neutralization by polyclonal anti-Ad serum was also achieved using both ligands, albeit less effectively than against mAb 9C12. Studies have shown that serum post-vaccination with Ad vectors typically contains antibodies against other viral proteins in addition to hexon (including fiber), which are less likely to be masked by hexon ligation (*39*). Our platform has the potential to shield the Ad virion from other potentially undesirable capsid interactors. For instance, display of DogCatcher-RBD blocked hFX-mediated transduction of SKOV-3 cells *in vitro*, presumably by inhibiting interaction between hFX and hexon (Fig. 4F). Shielding this interaction *in vivo* would be advantageous, since hFX enhances Ad5 transduction of hepatocytes, increasing liver sequestration and hepatotoxicity after systemic administration (*26*). A recent study has also reported an interaction between capsid-associated hexon of various serotypes and platelet factor 4 (PF4), which may contribute towards the development of very rare blood clots associated with some COVID-19 vaccines (*40*). Further studies will be required to assess the capability of our platform to inhibit interactions between hFX (or other host proteins) and hexon *in vivo* but this platform could represent an attractive solution to achieve capsid shielding. To date, Ad capsid shielding has predominantly involved coating virions with polymers, such as polyethylene glycol (PEG) or N-(2-hydroxypropyl)methacrylamide (HPMA) (*41*). Although effective in shielding, these polymer coatings have tended to impair vector transduction (*42*).

Capsid decoration with DogCatcher-RBD significantly enhanced boosting of humoral immunity against SARS-CoV-2 using Ad vaccine vectors. In homologous prime-boost regimens, decorating an Ad vector encoding spike with DogCatcher-RBD increased median anti-RBD antibody responses and pVNT titers >10 fold compared to the undecorated Ad (Fig. 6). Strikingly, a vector encoding an irrelevant antigen (GFP) but with RBD displayed on the Ad capsid also elicited significantly higher anti-RBD antibody responses compared to the undecorated Ad encoding spike, and median pVNT titers were >10-fold higher against the Wuhan strain and all variants of concern except delta (6.2-fold). Key to the enhanced humoral immunity achieved with our platform was increased boostability of antibody responses against capsid-displayed RBD. Fold-increase in anti-RBD antibody titers between prime and boost was ∼10-fold higher with RBD capsid display compared to the undecorated Ad (Fig. 6D). A similar observation was made after a heterologous regimen using RBD-decorated Ads to boost immunity following immunization with a conventional undecorated Ad encoding spike (Fig. 7A). In both experiments, boosting of humoral immunity using homologous prime-boost with undecorated Ad encoding spike was modest (∼2-5-fold increase in anti-RBD ELISA, Fig. 6D and 7B), and these data are comparable to previous reports using other Ad vectors encoding SARS-CoV-2 spike in mice (*43*). In trials of COVID-19 vaccines in humans, boosting of cellular and humoral immunity using homologous Ad prime-boost regimens was also modest, particularly when compared to heterologous regimens with mRNA vaccines or homologous mRNA regimens (*44-46*). Anti-vector immunity generated after a prime immunization may inhibit the ability to boost immune responses to encoded antigens using the same vector; anti-Ad neutralizing antibodies prevent vector transduction and subsequent transgene antigen expression required for antigen-specific immunity (*9*). In contrast, recombinant protein antigens displayed in particulate form on nanoparticles or VLPs (in adjuvant formulations) have been shown to generate robust humoral immunity, with highly efficient boosting in homologous prime-boost regimens (*47, 48*). Previous studies have suggested that particulate antigens generate potent antibody responses through a variety of mechanisms, including B cell receptor crosslinking (*49*) and prolonged retention in draining lymph nodes (*50*). It seems likely that responses against capsid-decorated RBD are induced by similar mechanisms, particularly since Ad particles have an optimum size for lymph node entry and retention (*51*), although future studies will be required to elucidate these mechanisms further.

While capsid antigen decoration induced potent humoral immunity, encoding of antigen sequences in the vector genome was required to generate potent cellular immunity (Fig. 4F). This observation is unsurprising, considering Ad vaccine vectors are known to induce particularly strong T cell immunity, especially CD8^+^ T cell responses (*6, 52*). Importantly, capsid decoration did not impair T cell responses to encoded antigens (Fig. 4G, 6F and 6G), and in fact a heterologous prime-boost regimen using capsid-decorated vectors at boost improved cellular immunity compared to a homologous regimen (Fig 7C and 7D). The latter observation suggests that our capsid shielding technology applied at boost may have, at least in part, circumvented anti-vector immunity generated after a priming immunization, as observed *in vitro* (Fig. 3 and Fig. 4). In support of this hypothesis, median SARS-CoV-2 spike-specific IFNγ ELISpot responses were comparable between homologous prime-boost regimens using decorated and undecorated vectors encoding spike in both experiments (Fig. 6G and 7C) but were ∼2 fold higher in the heterologous regimen (Fig. 7C).

Our capsid display platform offers a substantial improvement over conventional Ad vaccine technology for induction of both cellular and humoral immunity. Taken together, the data presented here support a concept for optimizing concomitant humoral and cellular immunogenicity, whereby antigenic targets for antibody induction are displayed on the capsid surface, while potent T cell epitopes are encoded in the vector genome. Such a concept would enable conserved components of target pathogens that induce potent T cell responses (such as nucleocapsid (*53*) or ORF1ab (*54*) of SARS-CoV-2) to be encoded in the vector genome, while targets of neutralizing humoral immunity such as RBD (often more heterogeneous in sequence) can be displayed on the capsid surface. Surface-displayed targets may even be replaced with different versions in response to the emergence of new pathogen variants. Indeed, a recent study has suggested that memory T cells recognizing epitopes within the SARS-CoV-2 replication-transcription complex that are shared with seasonal human coronaviruses may have protected against SARS-CoV-2 infection in a cohort of healthcare workers who remained seronegative (*55*). Further investigation will be required to comprehensively determine limitations for capsid display in terms of size and structure of ligands, but similar sized receptor-binding domains of other viral proteins, including influenza hemagglutinin (∼25 kDa), have been expressed independently as recombinant proteins and displayed on VLPs (*56*) implying that this could be a generalizable concept.

Capsid decoration using our protein superglue technology is simple, requiring only co-incubation of spontaneously and irreversibly reacting components with no chemical modification required. A similar conjugation process has already been scaled under good manufacturing practice (GMP) during development of a VLP-based SARS-CoV-2 vaccine currently in Phase I/II clinical trials (*19*). The ability of our adenovirus-based platform to induce both robust cellular and humoral immunity and to enhance efficacy of multi-shot regimens could be advantageous for applications beyond prophylactic vaccines, including therapeutic vaccines against chronic viral pathogens and cancer. Methods of rapid and customizable covalent decoration of Ad capsids could also be utilized for development of personalized therapies. In prophylactic settings, adenovirus capsid decoration could be utilized in the design of pan-coronavirus and pan-influenza vaccines; combining broad and conserved T cell immunity from encoded antigens with exchangeable capsid ligands delivering potent neutralizing humoral immunity.

## MATERIALS AND METHODS

### Construction of Ad-DogTag vectors

An expression construct, consisting of the cytomegalovirus immediate early promoter (CMVp) containing tetracycline operator sequences driving expression of enhanced green fluorescent protein (GFP), was cloned into the shuttle vector pENTR4 (Invitrogen). The CMVp GFP expression construct was inserted into an E1/E3-deleted (replication-defective) molecular clone of Ad5 at the E1 locus using Invitrogen Gateway site-specific recombination technology. Bacterial artificial chromosome (BAC) sequences from pBELOBAC11 (NEB) were amplified using forward (5’-TTAATTAAcgtcgaccaattctcatg) and reverse (5’-TTAATTAAgtcgacagcgacacacttg) primers to introduce *PacI* sites at either end of the BAC cassette. The entire Ad5(GFP) genome was subsequently cloned into the BAC with PacI, to generate pBAC-Ad5(GFP). DogTag (DIPATYEFTDGKHYITNEPIPPK) or SpyTag (AHIVMVDAYKPTK) sequences flanked by GSGGSG linkers (Fig. 1B) were inserted into hexon HVR1, HVR2 or HVR5 loops through BAC GalK recombineering, using the *GalK* gene for both positive and negative selection as described previously (*57*). Insertion sites in the hexon are shown in Fig. 1B; insertions at similar sites have been described previously (*29, 58, 59*). In this study, ‘DogCatcher’ refers to the sequence described as ‘R2CatcherB’ in Keeble *et al* (*21*). Recombinant vectors expressing DogCatcher-NANP18 and SARS-CoV-2 spike (residues 1-1208 Wuhan strain, codon-optimized for mammalian expression and including stabilizing mutations K986P and V987P and mutation of the furin cleavage site 682-GSAS-685 (*60*)) were generated through cloning of gene constructs into pENTR4.CMVp and subsequent Gateway-mediated insertion into the Ad5 E1 locus.

### Rescue and purification of recombinant adenoviruses

BAC DNA from recombinant molecular clones was linearized with PacI to release left and right viral inverted terminal repeats (ITRs). Linearized DNA was transfected (Lipofectamine 2000, Invitrogen) into E1-complementing Human Embryonic Kidney (HEK) 293 cell lines; either 293A cells (Invitrogen) for Ad vectors expressing GFP and DogCatcher-NANP18, or 293TREX cells (Invitrogen) for Ad vectors expressing SARS-CoV-2 spike. After cytopathic effect (CPE) was observed, the cells were harvested, subjected to three cycles of freeze-thaw, and the virus amplified further. Upon infection of 10 × 150 cm^2^ flasks, virus was harvested from infected cells after 48 h and purified by CsCl gradient ultracentrifugation according to standard protocols. Purified virus was dialysed against sucrose storage buffer (10 mM Tris-HCl, 7.5% w/v sucrose, pH 7.8) and stored at −80°C.

### Titration of recombinant adenoviruses

Infectious titer of vector preparations was assessed by single cell infectivity assay on HEK293A cells or 293TREX cells. For vectors expressing GFP, infected HEK293A cells were visualized and enumerated directly by fluorescent microscopy. For vectors without a fluorescent marker, cells (either 293A or 293TREX) were immunostained for expression of hexon. Serial dilutions of virus in complete media (Dulbecco’s Modified Eagles Medium, plus 1× GlutaMAX and 10% v/v fetal bovine serum) were performed in 96-well deep well plates. Two or three serial dilutions were performed per virus, and 50 µL of each dilution (10^3^ to 10^10^ dilution) was added per well of a seeded 96-well plate at 80-90% confluency. Plates were incubated for 48 h at 37°C, 5% v/v CO_2_. For titration of GFP-expressing vectors, single GFP-positive cells were enumerated by fluorescence microscopy, and an infectious titer was calculated in infectious units (ifu) per mL. For hexon immunostaining, media was aspirated from the cell monolayer and cells were fixed with ice-cold methanol. Plates were then washed in phosphate buffered saline (PBS 1X, Gibco) before blocking for 1 h with 3% w/v low-fat milk (Marvel). Mouse monoclonal anti-hexon antibody (B025/AD51, Thermo-Fisher) was added at 1:1000 dilution in 1% w/v milk in PBS and incubated for 1 h at 25°C. Cells were washed with 1% w/v milk in PBS prior to addition of a secondary goat anti-mouse alkaline phosphatase (ALP) conjugated antibody (STAR117A, BioRad) at 1:1000 dilution in 1% w/v milk in PBS. After a further 1 h incubation, plates were washed with PBS prior to development. To develop, 100 µL of freshly prepared SIGMAfast BCIP/NBT solution (Sigma-Aldrich) was added to each well and plates incubated at 25°C prior to the appearance of dark stained foci, representing single infected cells. For CsCl-purified vector preparations, viral particle count was estimated by spectrophotometric absorption at 260 nm as described previously (*12, 61*). P:I ratios were calculated by dividing the number of viral particles (vp) per mL with the number of infectious units (ifu) per mL.

### Protein production and purification

DNA sequences for expression of DogCatcher, DogCatcher-NANP_9_, DogCatcher-NANP_18_ and SpyCatcher were cloned into expression plasmid pET45(+) (EMD Millipore) for protein production in BL21(DE3) *E. coli* (NEB). Recombinant proteins were purified using Ni-NTA affinity resin (Qiagen) according to standard protocols (*21*), dialysed into PBS, and stored at -80°C.

DNA sequences for expression of DogCatcher fused to SARS-CoV-2 spike receptor binding domain (Wuhan strain, residues 319-532) (DogCatcher-RBD), were cloned into mammalian protein expression plasmid pcDNA3.4. To facilitate secretion, the Igκ-leader sequence METDTLLLWVLLLWVPGSTGD was introduced at the N-terminus of the fusion protein, and a C-terminal C-tag (EPEA) was added to enable affinity purification. DogCatcher-S-RBD was expressed in suspension ExpiCHO-S cells (Thermo Fisher); protein was harvested from culture supernatant, affinity purified using C-tag affinity resin (Thermo Fisher) using an AKTA chromatography system (GE Healthcare), and dialysed into Tris-buffered saline (TBS) pH 7.4.

Proteins for ELISA assays, RBD-SnJr (RBD with SnoopTagJr fused at the C-terminus) and spike-SnJr (SARS-CoV-2 spike 1-1208, with stabilizing proline mutations and furin cleavage site mutation (*60*), a C-terminal gp160 trimerization domain and SnoopTagJr) were expressed in suspension Expi293F (Thermo Fisher) and ExpiCHO-S cells respectively and purified as described above for DogCatcher-RBD.

### Coupling reactions

For *in vitro* assays, coupling reactions between DogCatcher-fused protein ligands and DogTag inserted into the Ad capsid were performed by co-incubation of spontaneously reacting components in a total volume of 25 μL, with individual components at concentrations described in the figure legends. Reactions were incubated for 16 h at 4°C.

Ligand-decorated vector batches for the NANP vaccine studies were prepared by co-incubating 1.9E+12 viral particles of Ad5(GFP)-HVR5-DogTag with 35 μM DogCatcher-NANP18 for 16 h at 4°C. To remove excess ligand, coupled vectors were dialysed into sucrose storage buffer using SpectraPor dialysis cassettes with 300 kDa molecular weight cut-off (MWCO). Dialysis decreased the excess ligand by at least 10-fold, as measured by Coomassie-stained SDS-PAGE. Ligand coverage by SDS-PAGE was >90% hexon (comparable to Fig. 3A).

Ligand-decorated vector batches for SARS-CoV-2 vaccine studies and electron microscopy were prepared by co-incubating 9E+11 viral particles of Ad5-HVR5-DogTag encoding either GFP or SARS-CoV-2 spike with 6 μM DogCatcher-RBD for 16 h at 4°C. To remove excess ligand, coupled vectors were dialysed into sucrose storage buffer using SpectraPor dialysis cassettes (300 kDa MWCO). Dialysis decreased excess ligand by at least 20-fold as measured by Coomassie-stained SDS-PAGE. Ligand coverage by SDS-PAGE was ∼60% hexon (comparable to Fig. 4B).

For all decorated vaccine vectors, residual excess ligand post-dialysis was factored into effective antigen dose calculations. Ligand-decorated vectors were stored at -80°C, endotoxin tested, and infectious titration of stored batches was repeated prior to immunization.

### Assessment of coupling efficiency by SDS-PAGE

Coupling reactions were performed as described above and stopped by addition of SDS loading buffer (Bio-Rad, 31.5 mM Tris-HCl, pH 6.8, 10% glycerol, 1% SDS, 0.005% Bromophenol Blue, 300 mM dithiothreitol). Samples were boiled at 95°C for 5 min and loaded on SDS-PAGE (NuPAGE 4-12% Bis-Tris, Invitrogen). Efficiency of ligand coupling to the Ad capsid (% total hexon protein coupled) was assessed by direct gel shift assays. Proteins were resolved by SDS-PAGE (200 V, 55 min) and visualized by Coomassie staining [16 h staining with Quick Coomassie (Generon), destained with water]. Coupling efficiency was assessed by comparing band intensities of unconjugated hexon-Tag in ligand decorated samples to undecorated (control) samples using Image J:

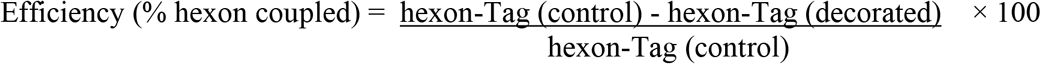

### Anti-vector antibody neutralization assay

For assessment of vector neutralization by potent neutralising mouse monoclonal antibody (mAb) 9C12 (*24*) (Developmental Studies Hybridoma Bank, University of Iowa), Ad5 vectors expressing GFP were incubated with serially diluted mAb 9C12 antibody at a 1:1 ratio in complete media for 1 h at 37°C. The Ad/9C12 mix was then added to an 80% confluent monolayer of HEK293A cells in a 96-well plate (cells were infected at a multiplicity of infection of 200 ifu/cell). Cells were incubated with the Ad/9C12 mix for 2 h at 37°C 5% v/v CO_2_, before the mix was replaced with fresh media and the plates returned to 37°C 5% v/v CO_2_ for a further 24 h. After 24 h, GFP expression was used as a readout of vector infectivity; bulk fluorescence was measured on a fluorimeter (Tecan) using an excitation wavelength of 395 nm and emission wavelength of 509 nm.

For assessment of vector neutralization by serum containing anti-adenovirus antibodies, serum samples were obtained by immunizing C57BL/6 mice with 1E+8 ifu of an Ad5 vector expressing ovalbumin (vector had an unmodified hexon). Serum was harvested two-weeks post immunization, stored at -80°C, and then serially diluted for the neutralization assay (two-fold dilutions were prepared from 1:4 to 1:512 in complete media, to give a final range of 1:8 to 1:1024 on cell monolayers). Diluted serum was incubated with Ad5(GFP) vectors, the mix incubated on HEK293 cells, and bulk GFP fluorescence read 24 h later as described above.

### Assessment of coagulation Factor X-mediated vector transduction of SKOV3 cells

SKOV3 cells (human ovary adenocarcinoma) were obtained from Public Health England and cultured in McCoy’s 5a media with 2 mM Glutamine and 15% v/v fetal bovine serum (complete McCoy’s). For the assay, GFP-expressing Ad vectors were serially diluted (1:10 to 1:10^7^) in serum-free media. Human coagulation Factor X (hFX) (Haematologic Technologies) was added to diluted vectors at a final concentration of 8 μg/mL. Ad/hFX mixes were added to monolayers of SKOV3 cells in 96-well plates and incubated with cells for 2.5 h at 37°C and 5% v/v CO_2_. After 2 h, Ad/hFX mixtures were replaced with complete McCoy’s media, and plates incubated at 37°C, 5% v/v CO_2_ for a further 48h. Infectivity was assessed after 48h by enumeration of GFP-positive foci as described above.

### Mouse immunizations

All mouse procedures were performed in accordance with the terms of the UK Animals (Scientific Procedures) Act Project Licence (PA7D20B85) and approved by the Oxford University Ethical Review Body. Female BALB/c mice (age 6-8 weeks, Envigo), housed in specific-pathogen free environments, were immunized intramuscularly by injection of 50 μL of vaccine formulated in endotoxin-free PBS (Gibco) into both hind limbs of each animal (100 μL total). Doses of adenoviral vectors and recombinant proteins are described in figure legends. Protein vaccines were administered in combination with Alhydrogel (Invivogen) at a 1:9 v/v ratio of adjuvant to antigen. Endotoxin dose was < 3 EU per mouse in all studies. Experiments were performed at Biomedical Services, University of Oxford.

### Ex-vivo IFN-gamma ELISpot

Overnight spleen *ex vivo* interferon-gamma (IFN-γ) ELISpot was performed according to standard protocols as described previously (*62*). To measure antigen-specific responses, cells were re-stimulated with peptides for 18–20 h. To measure T cell responses to GFP, peptide CD8^+^ T cell epitope EGFP_200-208_ (*63*) was added at a final concentration of 5 μg/mL. To measure T cell responses to SARS-CoV-2 spike receptor binding domain (RBD), cells were stimulated with a peptide pool of 15-mer peptides with an 11 amino acid overlap spanning the length of the RBD region (P0DTC2 319-541, PepMix™ SARS-CoV-2 S-RBD, JPT Peptide Technologies). Pooled peptides were added at a final concentration of 0.5 μg/mL for each peptide. To measure T cell responses to SARS-CoV-2 spike protein, cells were stimulated with two peptide pools of 15-mer peptides with an 11 amino acid overlap spanning the entire length of the spike protein (PepMix™ SARS-CoV-2 spike glycoprotein, JPT Peptide Technologies). Pooled peptides were added at a final concentration of 0.5 μg/mL for each peptide. Spot forming cells (SFC) were measured using an automated ELISpot reader system (AID).

### IgG ELISA

IgG endpoint ELISA was performed as described previously (*64*). Plates were coated with either recombinant GFP protein (Millipore) at 1 μg/mL to measure GFP-specific responses, recombinant DogCatcher-NANP18 protein at 2 μg/mL to measure DogCatcher-NANP18-specific responses, or either recombinant RBD-SnJr or spike-SnJr at 2 μg/mL to measure SARS-CoV-2 specific responses. To generate endpoint titers, sera from mice immunized with Ad5 vectors expressing an irrelevant antigen were used as a negative control.

### SARS-CoV-2 pseudovirus neutralization (pVNT) assay

Pseudotyped HIV-1 viruses incorporating the SARS-CoV-2 full-length spike (Wuhan strain or B.1.1.7, B.1.617.2 or B.1.351 variants of concern) were generated and SARS-CoV-2 pVNT assays performed as previously described using Hela cells stably expressing ACE2 as target cells (*28*). Serum was heat-inactivated at 56°C for 30 min prior to use in the assay.

### CryoEM data collection and image processing

CryoEM analysis was performed by NanoImaging Services, San Diego, USA. A 3 μL drop of sample Ad-DogTag (Control) and sample Ad-DogTag:DogCatcher-RBD were applied to a 1/2 Cu-mesh C-flat grid that had been plasma-cleaned for 10 s using a 25% v/v O_2_ / 75% v/v Ar mixture in a Solarus 950 Plasma Cleaner (Gatan). Grids were manually plunge frozen in liquid ethane. Data collection was carried out using a Thermo Fisher Scientific Glacios Cryo Transmission Electron Microscope operated at 200 kV and equipped with a Falcon 3 direct electron detector. Automated data-collection was performed with Leginon software (*65*) at a nominal magnification of 28,000×, corresponding to a pixel size of 5.19 Å. A total of 845 and 617 movies were recorded for samples Ad-DogTag and Ad-DogTag:DogCatcher-RBD, respectively, using a nominal defocus range of −2.4 to −5.6 μm. Exposures were fractionated into 19 frames with an exposure rate of 2.6 e^−^/pixel/s and total exposure of 10 e^−^/Å^2^.

For both samples, motion correction and contrast transfer function (CTF) estimation were performed using cryoSPARC3.1 (*66*). Using the Gaussian blob picker in cryoSPARC3.1, a total of 28,371 particles were picked for sample Ad-DogTag:DogCatcher-RBD and 91,920 particles were picked for sample Ad-DogTag. These particles were extracted from the cryoEM images and subjected to reference-free 2D classification in the cryoSPARC3.1. The best 28,307 particles for sample Ad-DogTag:DogCatcher-RBD and all particles picked for sample Ad-DogTag were used for the subsequent 3D reconstruction. An initial model was generated ab initio from all selected particles for sample Ad-DogTag:DogCatcher-RBD and used during one round of homogeneous refinement with icosahedral symmetry imposed in cryoSPARC3.1. The final 3D reconstruction of Ad-DogTag:DogCatcher-RBD was low-pass filtered to 30 Å and used as a starting volume for the 3D reconstruction of sample Ad-DogTag during one round of homogeneous refinement with icosahedral symmetry imposed in cryoSPARC3.1. The resolution of the final 3D reconstructions was determined by the Fourier shell correlation (FSC) between two independent half maps, which was 10.5 Å at FSC = 0.143 for both samples. Maps are visualized using Chimera (*67*).

### Statistics

Statistical analyses were performed in GraphPad Prism 9. Comparisons between multiple groups were performed by Kruskal-Wallis with Dunn’s test for multiple comparisons. Pair-wise comparisons (two groups only) were performed by Mann-Whitney test. Statistical significance indicated as follows: *p<0.05, **p<0.01, ***p<0.001, ****p<0.0001, ns not significant.

## Supplementary Materials

Fig. S1. DogTag/DogCatcher covalent coupling

Fig. S2. Yield of adenovirus vectors displaying SpyTag or DogTag at hexon HVR 1, 2 or 5.

Fig. S3. SpyTag is poorly reactive with SpyCatcher following insertion into adenovirus hexon HVR loops, with highly decorated virions losing infectivity.

Fig. S4. CryoEM; fitting RBD structure into density maps of Ad-Tag:Catcher-RBD

Fig. S5. Serum IgG responses against SARS-CoV-2 spike ectodomain

## Acknowledgments

The authors would like to thank Genevieve Labbe, Sophie Porret, Laura Bilbé, Rebecca Dabbs and Jing Jin for helpful advice and assistance. We thank Viv Clark, Heather Chandler, Stephen Laird, Douglas Passos and Luke Harris for animal husbandry and technical assistance. We thank Sarah Dunn, Weili Zheng, and Mandy Janssen from NanoImaging Services, San Diego, USA for cryoEM services and helpful discussion.

## Author contributions

Conceptualization: MDJD, SJD, MH, SB

Methodology: MDJD, CG, KJD, MHM, SJD, MH, SB

Investigation: MDJD, LMR, LAHB, CG

Visualization: MDJD

Funding acquisition:

Resources: MH

Writing – original draft: MDJD

Writing – review & editing: MDJD, MH, SB

## Data and materials availability

The cryo-EM maps for samples Ad5-DogTag (Control) and Ad5-DogTag:DogCatcher-RBD were deposited in the EMDB with accession codes EMD-14371 and EMD-14370 respectively. All other data are available in the main text or the supplementary materials.

