## Supplementary Figures S1 - S5 for "Modular capsid decoration boosts adenovirus vaccine-induced humoral and cellular immunity against SARS-CoV-2"

### Supplementary Materials

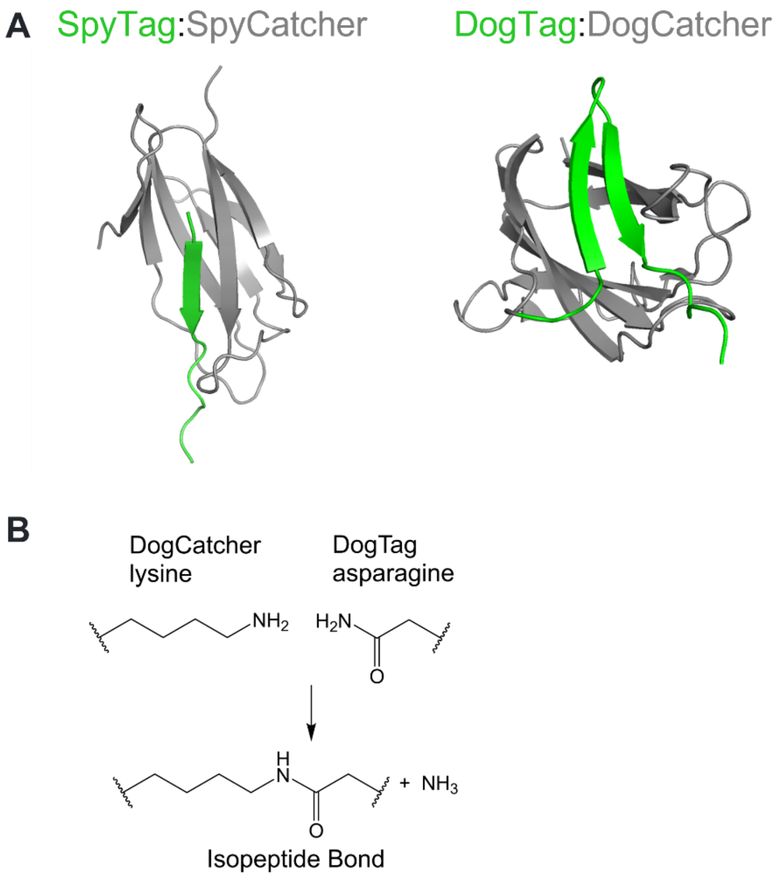


**Fig. S1. DogTag/DogCatcher covalent coupling**

(**A**) Comparison of SpyTag/SpyCatcher and DogTag/DogCatcher complexes. (Left) In the SpyTag/SpyCatcher complex, the tag is a linear single β-strand (model based on PDB structure 4MLI, SpyTag shown in green). (Right) In the DogTag/DogCatcher complex, the tag is a β-hairpin structure arranged from two β-strands (model based on PDB structure 2WW8 of *Streptococcus Pneumoniae* RrgA adhesin domain 4, DogTag shown in green). (**B**) Chemistry of isopeptide bond formation between a lysine sidechain of DogCatcher and an asparagine sidechain of DogTag via a spontaneous transamidation reaction.

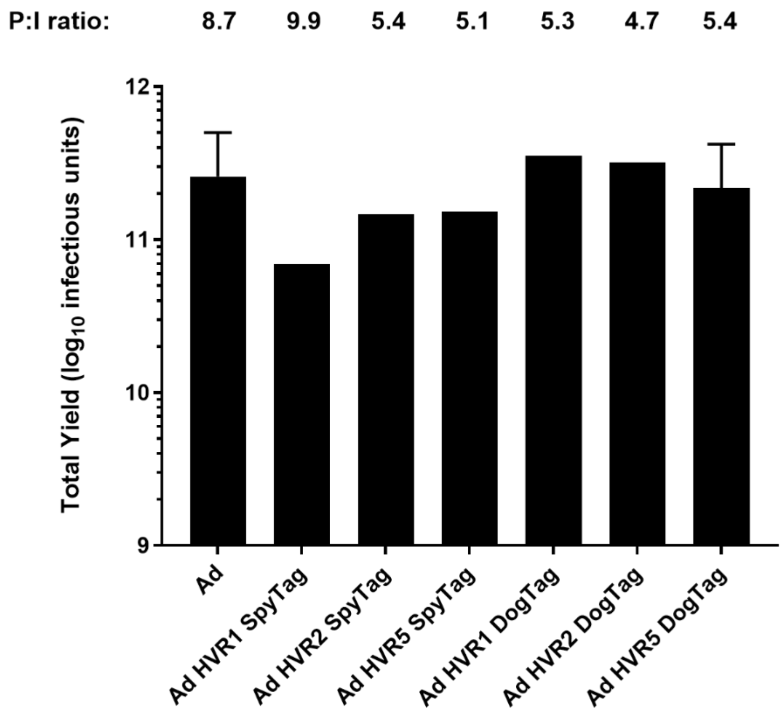


**Fig. S2. Yield of adenovirus vectors displaying SpyTag or DogTag at hexon HVR1, HVR2, or HVR5.**

Yield comparison of GFP expressing Ad vector preparations displaying either SpyTag or DogTag on hexon HVR1, 2 or 5. Data show infectious yield from 1500 $cm^2$ 293A cells, n=1 for all preparations except Ad5 (WT hexon) (n=3, see Fig. 2A) and Ad5 HVR5 DogTag (n=3, see Fig. 2A). Mean P:I ratios (ratio of total viral particles calculated by UV spectrophotometry to infectious units calculated by GFP focus assay) for vector batches are indicated above each bar.

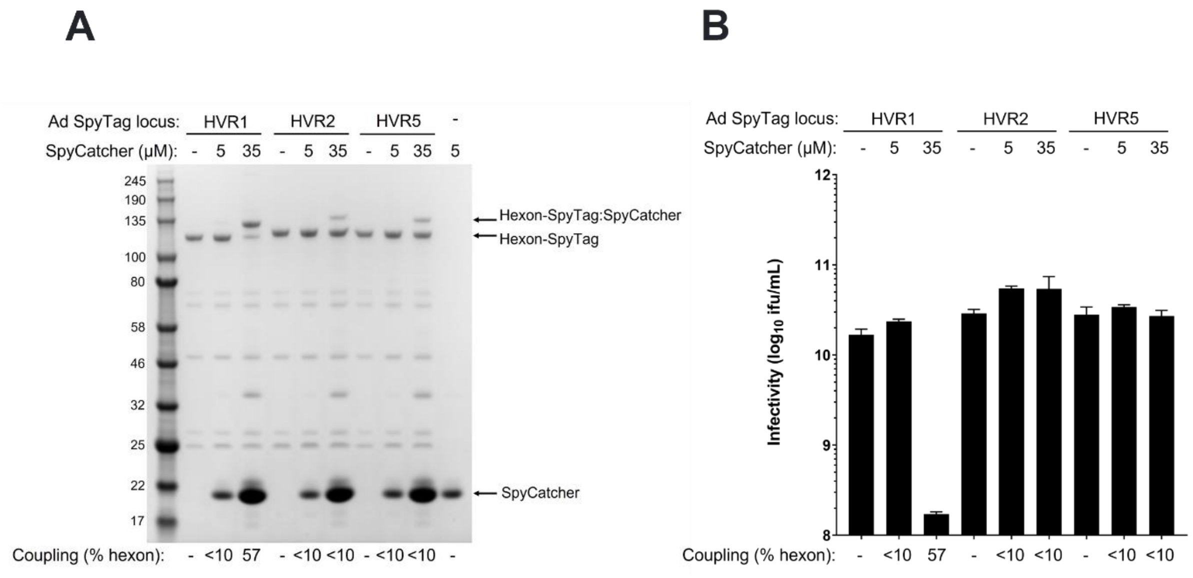


**Fig. S3. SpyTag is poorly reactive with SpyCatcher following insertion into adenovirus hexon HVR loops, with highly decorated virions losing infectivity.**

(**A**) SDS-PAGE and Coomassie staining analysis of Ad virions displaying SpyTag at HVR1, HVR2 or HVR5 (1E+10 viral particles) incubated with SpyCatcher (5 μM or 35 μM) at 4°C for 16 h. (**B**) Vector infectivity (GFP focus assay) performed on the samples from A (all vectors express encoded GFP). Data show mean + range of duplicate wells.

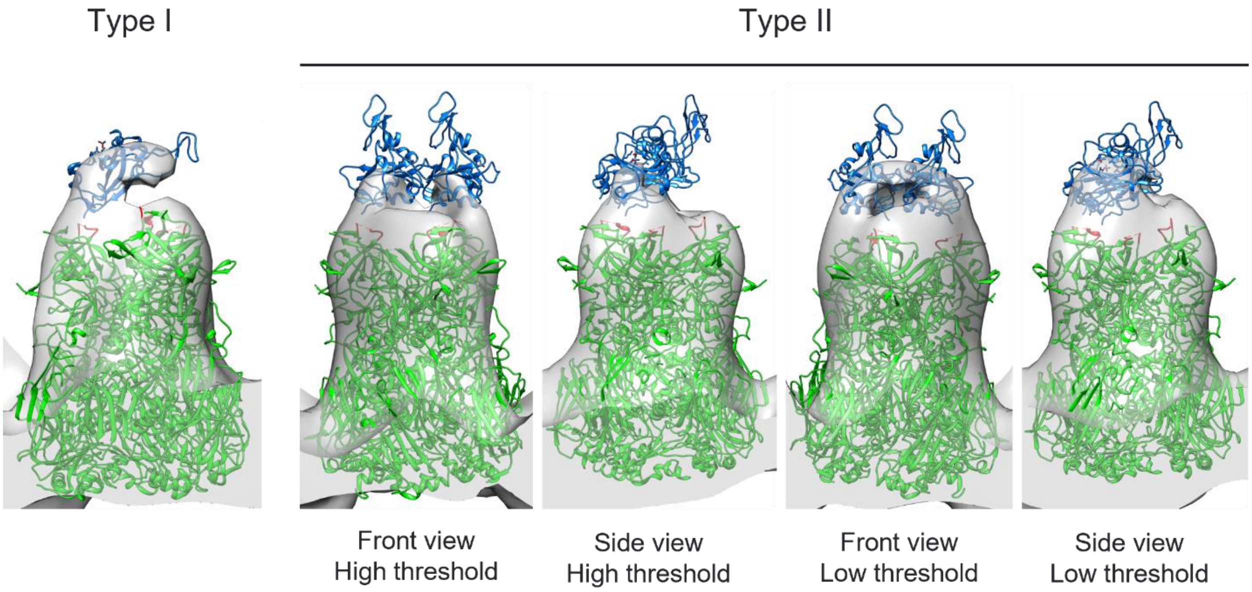


**Fig. S4. CryoEM; fitting RBD structure into density maps of Ad-DogTag:DogCatcher-RBD**

Structure of the SARS-CoV-2 spike receptor binding domain (RBD) (PDB ID J7VB) shown in blue was fitted into 3D density maps for Type I and Type II ligand coupling to hexon trimers on the surface of Ad-DogTag:DogCatcher-RBD from Fig. 5. Hexon trimer structure (PDB ID 6B1T) shown in green, with location of HVR5 loop (residues 270-280, site of DogTag insertion) shown in red.

**A**

| Group | Vaccine | Label | Dose | Adjuvant | Regimen |
|---|---|---|---|---|---|
| 1 | Ad(encoded Spike)-DogTag | Ad(S)-T | $10^8$ infectious units | - | D0 Prime D21 Boost |
| 2 | Ad(encoded Spike)-DogTag:DogCatcher-RBD | Ad(S)-T:C-RBD | $10^8$ infectious units | - | D0 Prime D21 Boost |
| 3 | Ad(encoded GFP)-DogTag:DogCatcher-RBD | Ad(GFP)-T:C-RBD | $10^8$ infectious units | - | D0 Prime D21 Boost |
| 4 | DogCatcher-RBD recombinant protein | C-RBD | 0.2 μg protein | Alhydrogel | D0 Prime D21 Boost |

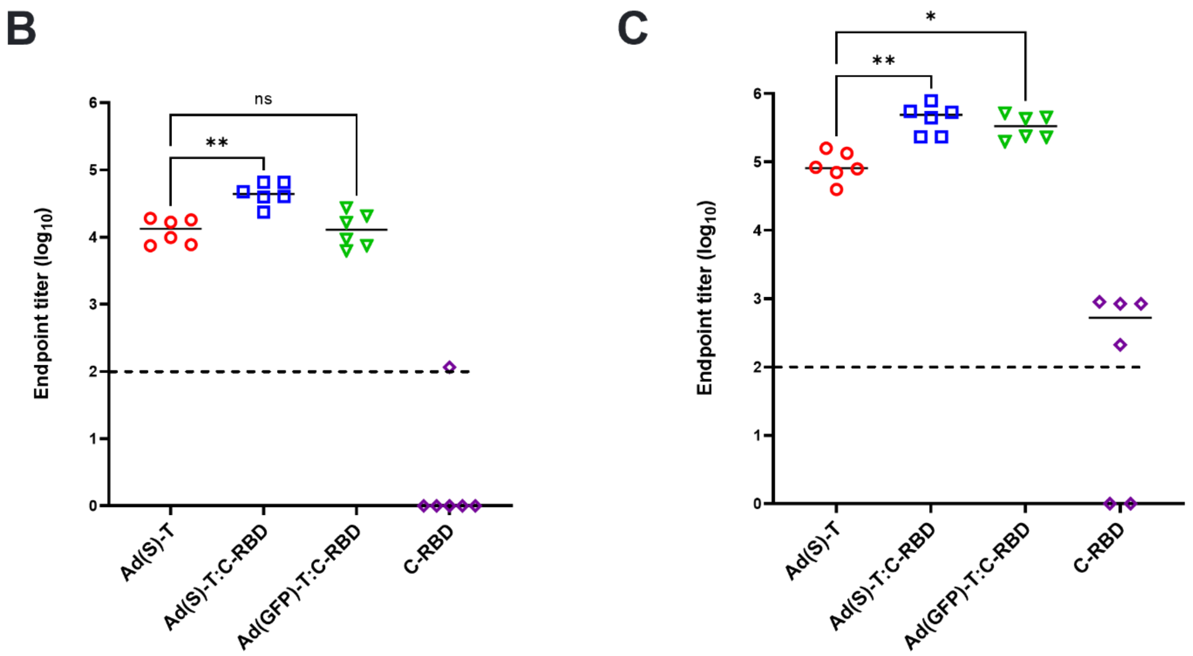


**Fig. S5. Serum IgG responses against SARS-CoV-2 spike ectodomain**

(**A**) Serum IgG responses against the full SARS-CoV-2 spike ectodomain were measured by endpoint ELISA in mice from the experiment shown in Fig. 6, groups of BALB/c mice were immunized intramuscularly as indicated. ELISA responses at (**B**) D20 and (**C**) D35 are shown. Median responses are shown as horizontal bars. Dashed line shows limit of detection.
